# Invasive *Aedes vittatus* mosquitoes in Jamaica: A looming public health concern

**DOI:** 10.1101/2025.04.10.648144

**Authors:** Simmoy A. A. Noble, Reneé L.M.N. Ali, Cameil F. Wilson-Clarke, Nadia K. Khouri, Douglas E. Norris, Simone L. Sandiford

## Abstract

*Aedes vittatus*, an emerging invasive mosquito of significant public health concern has slowly made its way onto the global radar. With a known geographical range in Africa and Asia, where it is a competent vector for several arboviruses, this mosquito has now been reported in the Americas. As the spread of this mosquito has been partly linked to transcontinental trade and travel, Jamaica, the largest English-speaking country in the Caribbean, which serves as a central hub for trade and transport throughout the region has been on alert since its identification in neighbouring Dominican Republic and Cuba.

Through ongoing surveillance efforts from January 2023 to October 2024, we report the detection of *Ae. vittatus* across six locations in four parishes in Jamaica. Both larvae and adults were collected from rural and urban areas in the country. Additionally, we present the first complete annotated mitochondrial genomes of two specimens of this invasive mosquito species. Phylogenetic analysis based on a 539 bp fragment of the *cytochrome c oxidase subunit I* gene extracted from the derived mitochondrial genomes of Jamaican *Ae. vittatus*, revealed clustering with specimens from Cuba, India and Nepal. This study highlights the benefits of routine surveillance and the power of molecular approaches to identify invasive species and their potential origins.

**Author summary:** Increased global movement is facilitating the spread of invasive mosquito species and the pathogens they carry, posing significant public health risks. Jamaica, a popular tourist destination with rising visitor numbers and extensive trade connections, is particularly vulnerable. The invasive Asian tiger mosquito *Aedes albopictus*, a vector of numerous viruses to humans, was reported in Jamaica in 2019, highlighting this issue. Another invasive species of concern, *Ae. vittatus*, primarily found in Asia and Africa, has been detected in nearby Cuba and the Dominican Republic. This raises alarms about the potential for disease outbreaks as *Ae. vittatus* can also transmit viruses such a dengue and chikungunya.

We report the detection of *Ae. vittatus* in Jamaica in samples collected from January 2023 to October 2024. Its presence in multiple widely distributed locations suggests it may have been in Jamaica undetected for some time. This adaptability of this mosquito to human-inhabited areas increases the risk of pathogen transmission to persons, underscoring the urgent need for enhanced surveillance by the Vector Control program at the Ministry of Health and Wellness. Given its potential to transmit viruses, *Ae. vittatus* is emerging as a serious public health concern that requires immediate attention and action.

## Introduction

Reports of invasive arboviral vectors and mosquito-borne diseases of public health and veterinary concern have steadily increased across the globe [1]. Human-mediated activities, including trade and tourism, are among the primary pathways through which invasive species are introduced into new habitats [2]. In our interconnected society, the seamless mobility of people and objects facilitated by advancement and growth in transportation have profound implications for the introduction of species to new habitats and the transmission of infectious diseases [3]. Additionally, ecosystems may be altered by invasive species and resource competition may result in the displacement of existing species [4].

Jamaica’s position as a prominent tourist hotspot makes it vulnerable to the introduction and dissemination of re-emerging vectors and infectious agents. The country has seen a significant rise in total visitor arrivals to the island, increasing from 1,535,165 in 2021 [5] to 4,181,740 in 2023 [6]. Jamaica also functions as a transportation hub for the Caribbean region and has longstanding preferential bilateral trade agreements with countries such as Barbados, Guyana and Trinidad and Tobago. Furthermore, since the early 2000s Jamaica has established key partnerships with Cuba [7] and the Dominican Republic [8] and increases in both exports and imports have been noted within the last two years.

In the last century, the Asian tiger mosquito *Aedes albopictus* is the most successful invasive mosquito [9]. Native to Asia, this mosquito has become ubiquitous in the Caribbean and the Americas, and was reported in Jamaica in 2019 [10]. The distribution of this invasive species was facilitated primarily through human trade, where eggs were unintentionally transported on old tyres and decorative plants. Adult mosquitoes were also inadvertently carried by public and private ground transit from highly infested regions [9].

Currently, there is a new and emerging vector in the Western Hemisphere, *Ae. vittatus*. A member of the subgenus *Fredwardsius, Ae. vittatus* is native to Asia and parts of Africa [11, 12]. This, however, is rapidly changing in the New World, as *Ae. vittatus* has been reported from both the Dominican Republic [13] and Cuba [14, 15]. Genetic analysis conducted on specimens from both countries using short sequence fragments of the mitochondrial *cytochrome c oxidase subunit I* (COI) gene suggests multiple introductions from Asia into the Caribbean [13, 14]. Increasingly, however, complete mitochondrial genomes (mitogenome) are being utilized to provide greater insights into the evolutionary histories of mosquitoes [16]. While mitogenomes of the highly invasive *Ae. albopictus* mosquito [17-20] in addition to other invasive vectors such as *Ae. vexans* [21], *Ae. japonicus* [22, 23], and *Ae. scapularis* [24] have all been reported, no such data currently exists for *Ae. vittatus*.

### Vector competence and habitats

*Aedes vittatus* is known to be a competent vector for several arboviruses, including dengue [25], chikungunya [26, 27], Zika [28], yellow fever [12] and West Nile viruses [29]. Factors such as climate change, urbanization, lack of effective vector control [30] and resilient eggs [31] have allowed this invasive mosquito to thrive in new environments. Adults have been captured from forest, savannah and barren land [32]. The ability of *Ae. vittatus* to survive in a wide range of habitats speaks to its high environmental plasticity and is reflected in oviposition habitats ranging from natural to artificial containers. In Africa, *Ae. vittatus* primarily lay eggs in rock holes [33] and larvae have been collected from tree holes, fresh fruit husk, and puddles. However, in peridomestic habitats *Ae. vittatus* is predominantly found in artificial containers such as tyres, bottles, cups and potted plants, demonstrating the urbanization of this species in Nigeria, India and Pakistan [12, 34]. In southern India, larvae were collected from cement tanks, cement cisterns and mud pots [35]. Similar to what has been seen with *Ae. albopictus*, displacement to new regions and destruction of natural habitats has forced *Ae. vittatus* to adapt to breeding in human-made containers [1].

Addressing the emerging threat of invasive mosquitoes like *Ae. vittatus* requires a comprehensive approach that combines traditional entomological approaches and molecular tools. These include routine surveillance and reference sequence data for the development of innovative vector control strategies. This study reports the first detection of *Ae. vittatus* in Jamaica and the complete mitochondrial genome of these mosquitoes as a reference for these collections.

## Method

### Study area

As part of an ongoing arboviral surveillance project, convenience sampling was utilized to collect mosquito specimens from January 2023 to October 2024. Locations in St. Andrew were in and around the university thus facilitating ease of collections. Sampling of rural sites in St. Ann, St. Elizabeth and Westmoreland coincided with community engagement projects in those areas at that time. Mona is a peri-urban neighbourhood in the southeastern parish of St. Andrew, located 8 km from the capital of Kingston (Fig 1). It is a former sugar plantation and is where The University of the West Indies and a fresh-water reservoir are situated. Location 1 was an area of mixed vegetation with domestic animals (horses, sheep and chickens) present nearby (Fig 2a). Location 2 was a private residential community surrounded by a heavily forested area (Fig 2b) and was approximately 990 meters from location 1. Location 3 was a lot with overgrown shrubs and trees with a single building adjacent to both residential and abandoned houses (Fig 2c) and was 410 meters from location 1 and 1,290 meters from location 2.

**Fig 1.**
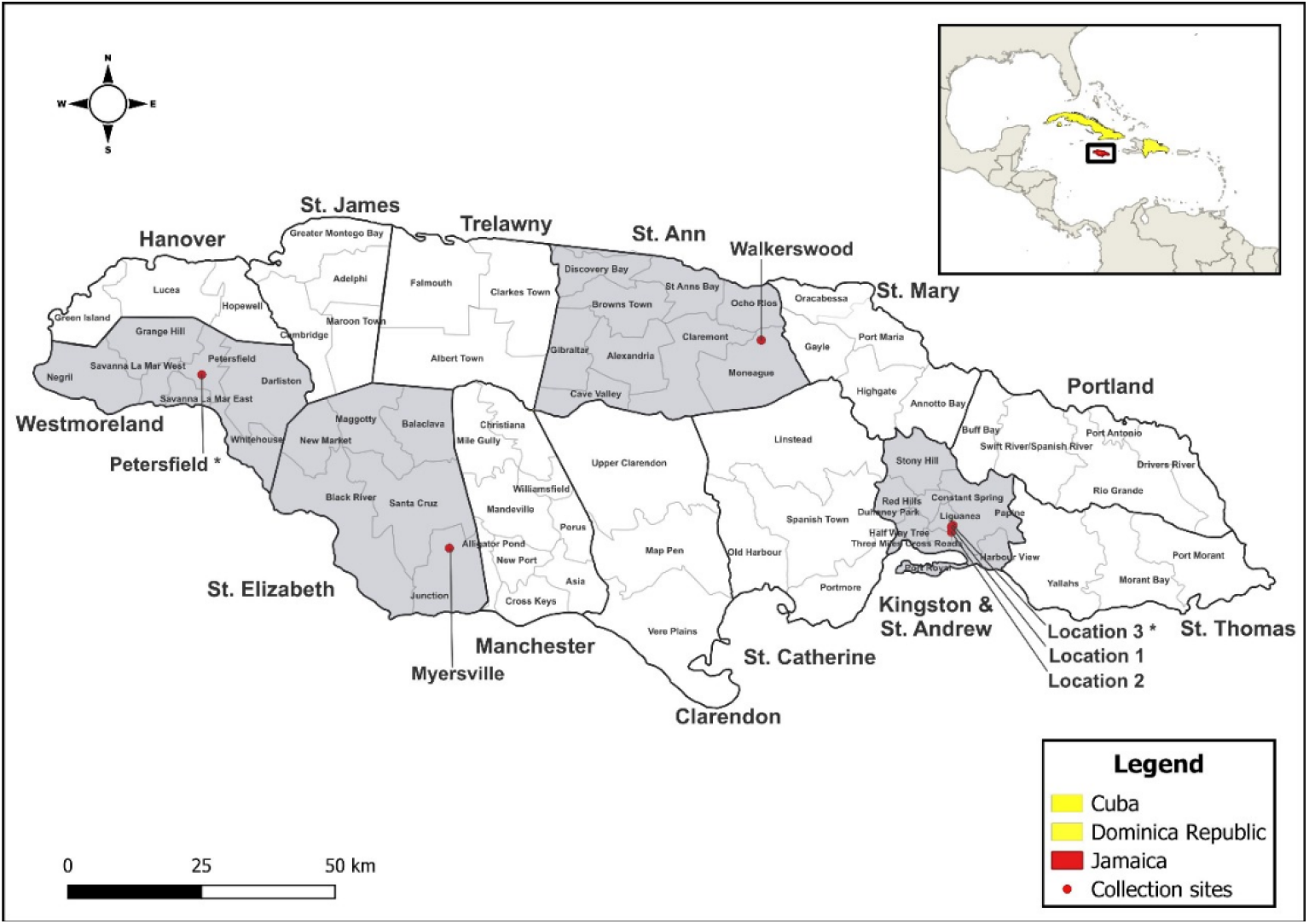
Map of Jamaica and sampled sites across the country. Collection sites are represented by the red dots and the asterisk indicates specimens that were used for mitogenome analysis. Yellow highlights countries in the Americas that have previously reported *Ae. vittatus*. Map was created using QGIS 3.34.11.

**Fig 2.**
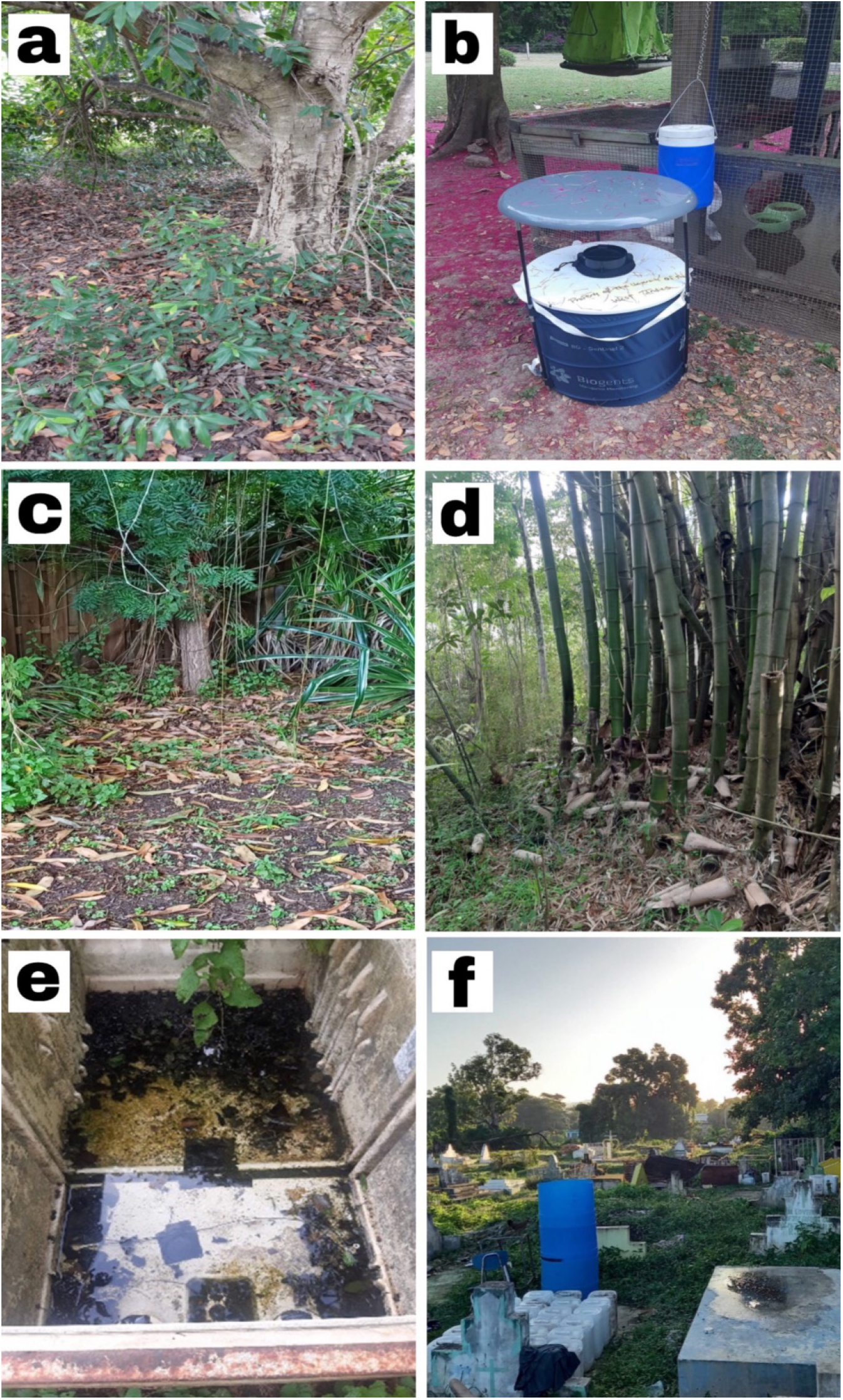
Locations where *Aedes vittatus* was identified. Location 1 Mona, St. Andrew. (b) Location 2 Mona, St. Andrew. (c) Location 3 Mona, St. Andrew. (d) Walkerswood, St. Ann. (e) Myersville, St. Elizabeth. (f) Petersfield, Westmoreland.

Walkerswood is a rural farming community in the parish of St. Ann in northern Jamaica (Fig 1). The collection site was a forested region at the back of a church, characterized by bamboo clusters and small to medium trees (Fig 2d). There was an indication of human activity at this site as many of the bamboo trees were deliberately cut down leaving the roots intact in the earth and presence of discarded containers (S1 Fig).

Myersville is a rural farming community in the parish of St. Elizabeth located in southwest Jamaica (Fig 1). Two locations were surveyed, the first a bushy area, with a large human-made water catchment area which provided a habitat for frogs and birds, and the second a forested area with large, discarded refrigerators positioned in the shade. The water inside the refrigerators was clear but contained decaying plant materials (Fig. 2e).

Petersfield is a rural sugarcane producing community in the parish of Westmoreland, which is situated 41 km southwest of Montego Bay, the second major city in Jamaica (Fig 1). The collection site was a cemetery, located behind a church and adjacent to a school and a heavily forested area (Fig 2f).

### Mosquito collection

Due to the proximity of the St. Andrew sites to The University of the West Indies, BG sentinel traps (Biogents, Germany) without BG chemical lure were set from 1400 to 1000 hr and baited with two pounds of dry ice to collect mosquitoes. At locations 1 and 3 in Mona the traps were positioned in shaded areas under trees whereas at location 2 it was positioned near an existing rabbit cage. Because of logistical challenges involved in conducting overnight trapping sessions at the rural locations, a Prokopack aspirator (John W. Hock, USA) was used to aspirate specimens from shrubs, during the hours of 1300 to 1500 hrs at the St. Ann and St. Elizabeth sites, and from 1100 to 1400 hrs at the Westmoreland sites. Adult specimens were transported to the laboratory on dry ice and stored at -80°C until sorting. Immature specimens were also collected from bamboo stems in St. Ann, discarded refrigerators in St. Elizabeth and a discarded plastic container in Westmoreland using disposable plastic pipettes. Larvae were reared to adults in site-collected water under standard laboratory conditions. All mosquitoes were morphologically identified using a Leica S9E stereomicroscope (Leica, Germany) and taxonomic keys to the species level [36, 37]. Select specimens in exceptional morphological condition were stored in tubes containing silica and shipped to Johns Hopkins Bloomberg School of Public Health for molecular analysis.

### DNA extraction, sequencing, mitogenome assembly and annotation

A modified extraction protocol was used to process single mosquito specimens which involved homogenization in a mixture of 98 μL of PK buffer (Applied Biosystems, Waltham, MA) with 2 μL of Proteinase K (Applied Biosystems, Waltham, MA) and incubation at 56°C as previously described [38, 39]. Following the manufacturer’s instructions (Qiagen DNeasy Blood and Tissue Kit, Hilden, Germany), DNA was extracted from the homogenate and quantified using the Qubit dsDNA assay kit (Thermo Fisher Scientific, Waltham, MA) prior to library construction and Illumina sequencing at the SeqCenter (Pittsburgh, USA). Using a genome skimming strategy, libraries were sequenced (2 x 150) to a depth of 13.3 million reads. Using the reference, *Ae. aegypti* (NC_035159.1) mitochondrial genome as the seed sequence and kmer set at 39, the *Ae. vittatus* mitogenomes were assembled in NOVOPlasty (RRID:SCR_017335) version 4.3.5 [40]. Automatic annotations using the invertebrate genetic code under default settings were identified in MITOchondrial genome annotation [41] on the publicly available Galaxy EU platform [42]. To match reference *Aedes* in the GenBank repository, start and stop codons were manually adjusted in Geneious Prime (RRID:SCR_010519) version 2025.0.3 (Biomatters, Auckland, New Zealand). Two (2) complete annotated mitochondrial genomes representing *Ae. vittatus* were submitted to GenBank.

### Phylogenetic analysis

The COI gene sequences extracted from the mitogenomes generated in this study were aligned with available reference *Ae. vittatus* and *Ae. aegypti* COI sequences from GenBank using MAFFT implemented in Geneious Prime (RRID:SCR_010519) version 2025.0.3 (Biomatters, Auckland, New Zealand). The alignment file was exported in nexus formatted and imported into the jModelTest (v2.1.10) software to determine best fit pair substitution model [43]. Using Bayesian Evolutionary Analysis by Sampling Trees (BEAST) 2 [44], Bayesian inference analysis was performed using three independent runs under default settings: using a 20 % burn in rate. FigTree v.1.4.4 was used to visualize trees (http://tree.bio.ed.ac.uk/software/figtree/).

## Results

### Collection of specimens

*Aedes vittatus* was identified in collections sorted for viral metagenomic analysis from the parishes of St. Andrew, St. Ann, St. Elizabeth and Westmoreland (Table 1). Although collected in multiple locations in 2023 and 2024, *Ae. vittatus* mosquitoes were of relatively low abundance compared to most other species except at the St. Elizabeth and Westmoreland sites where they comprised more than one quarter of the collected mosquitoes. Upon inspection, several cut bamboos contained both water and larvae in St Ann (Fig S1), however none of the larvae were *Ae. vittatus* (Table 1). Likewise, no *Ae. vittatus* larvae were found in the discarded container in Westmoreland (Table 1). Abandoned refrigerators at the St. Elizabeth served as a breeding site where immature *Ae. vittatus* were collected with *Ae. albopictus* larvae (Table 1).

**Table 1.**
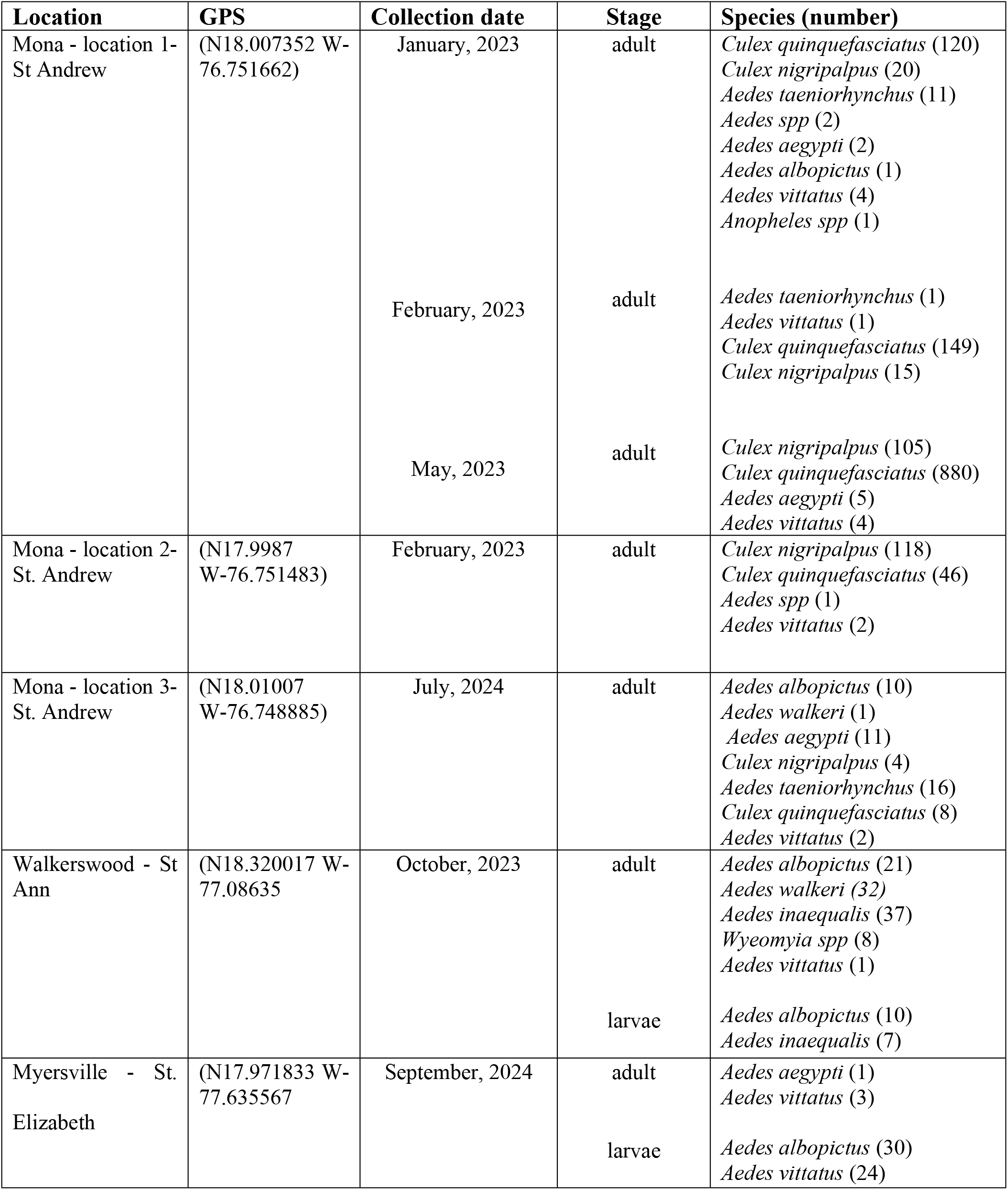
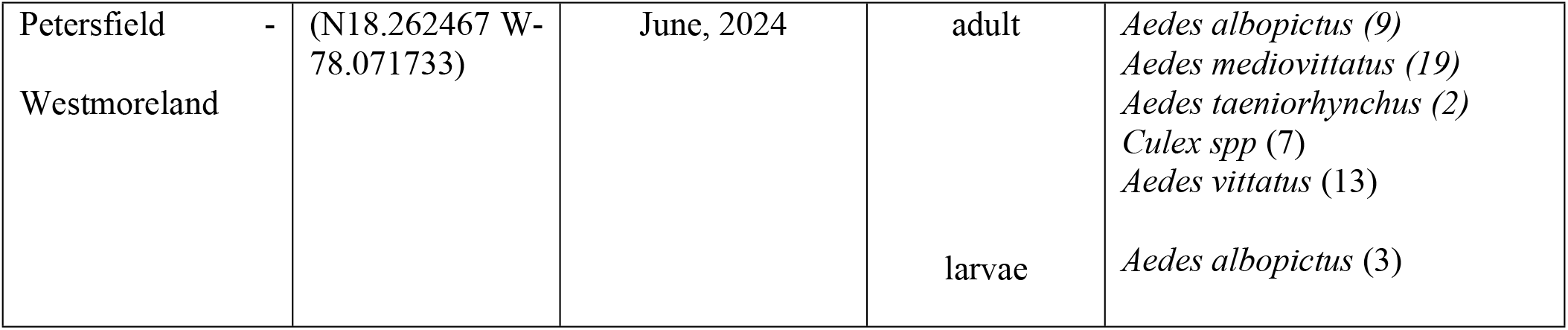
Mosquito species collected alongside *Aedes vittatus* in breeding and trap locations.

### Morphological and molecular identification of *Ae. vittatus* mosquitoes

Morphological identification of *Ae. vittatus* was confirmed by the presence of three pairs of small, spherical, silvery white spots on its scutum and white band at the base of the tibiae as seen in (Fig 3).

**Fig 3.**
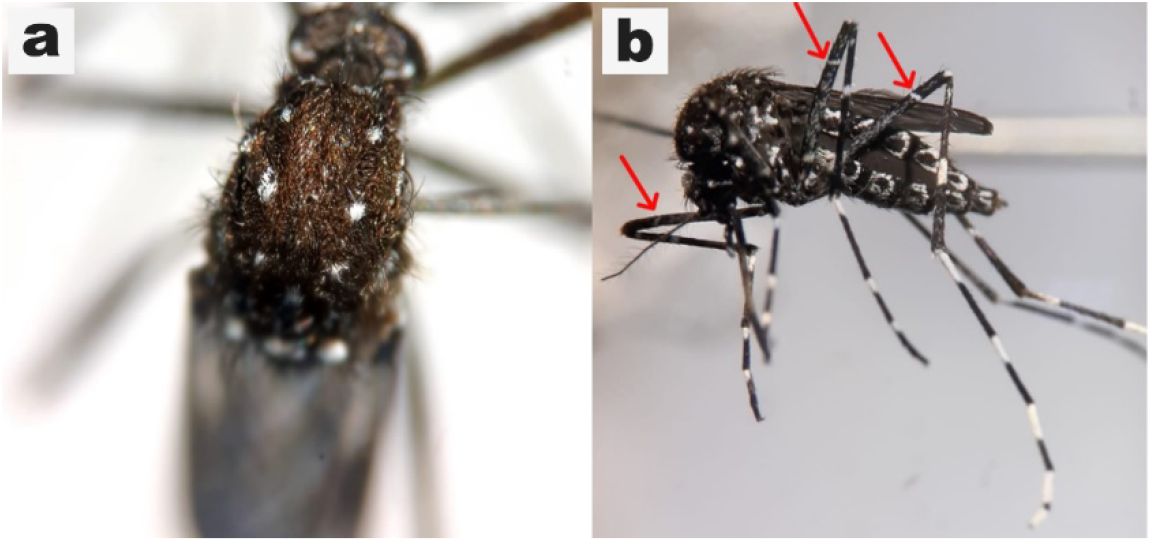
Morphological features of *Aedes vittatus*. **(a)**Scutum showing three pairs of slivery white spots. (**b)** All tibia with sub-basal white band.

To link morphological and molecular identification of *Ae. vittatus*, two high quality morphological specimens were randomly selected for Illumina sequencing and mitochondrial genome assembly. The mitogenomes generated in this study were 15,764 and 15,785 bp in length, with an average AT content of 79.3%. *Aedes vittatus* mitogenomes were comparable with other *Aedes* species with 37 genes: 13 protein coding genes, 22 transfer RNAs and 2 ribosomal RNA. The representative mitogenome map with annotated genes of *Ae. vittatus* is shown in (Fig 4).

**Fig 4.**
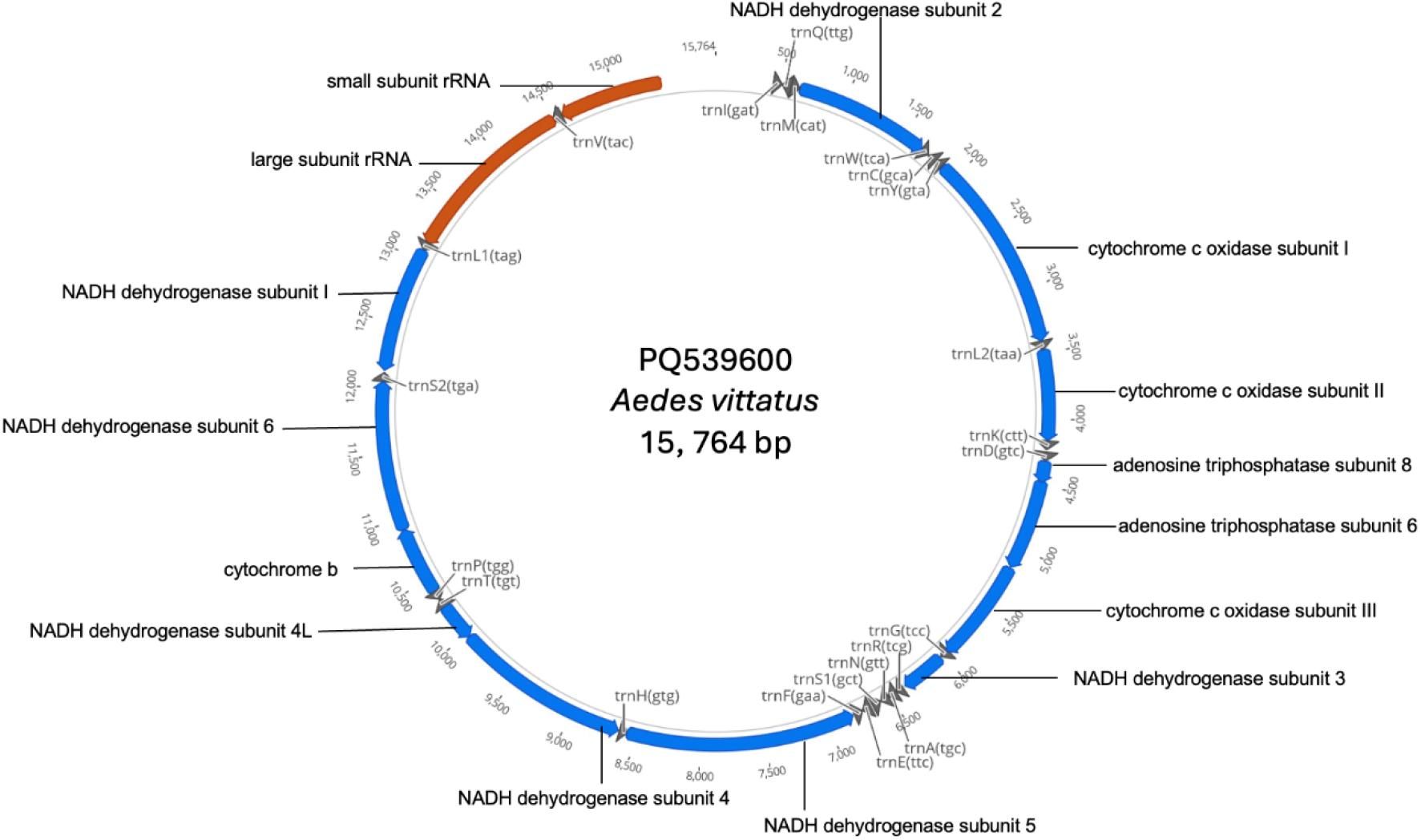
Representative mitochondrial genome map of *Ae. vittatus* from Mona, location 3 Jamaica. The blue, gray and red color blocks represent protein coding genes, transfer RNAs and ribosomal RNAs respectively.

Since no other *Ae. vittatus* mitogenome exists in the public domain, the COI gene segment (1531 bp in length) was extracted from each mitogenome and added to an alignment matrix (539 bp) with 15 *Ae. vittatus* COI sequences available from the GenBank repository. Results from the 539 bp COI fragment BLAST analysis showed that PQ539599 from Petersfield shared 99.63% identity with OL331077, which was obtained from a dengue-endemic region of Nepal [45]. In contrast, PQ539600 from Mona, location 3 shared 100% identity with MT519730 from Cuba [14]. Bayesian inferences resulted in the Caribbean, Southeast Asia and Africa COI sequences separating into 3 main clades: with the Jamaican *Ae. vittatus* COI sequences clustering with sequences of *Ae. vittatus* collected in Cuba, India and Nepal (Fig 5).

**Fig 5.**
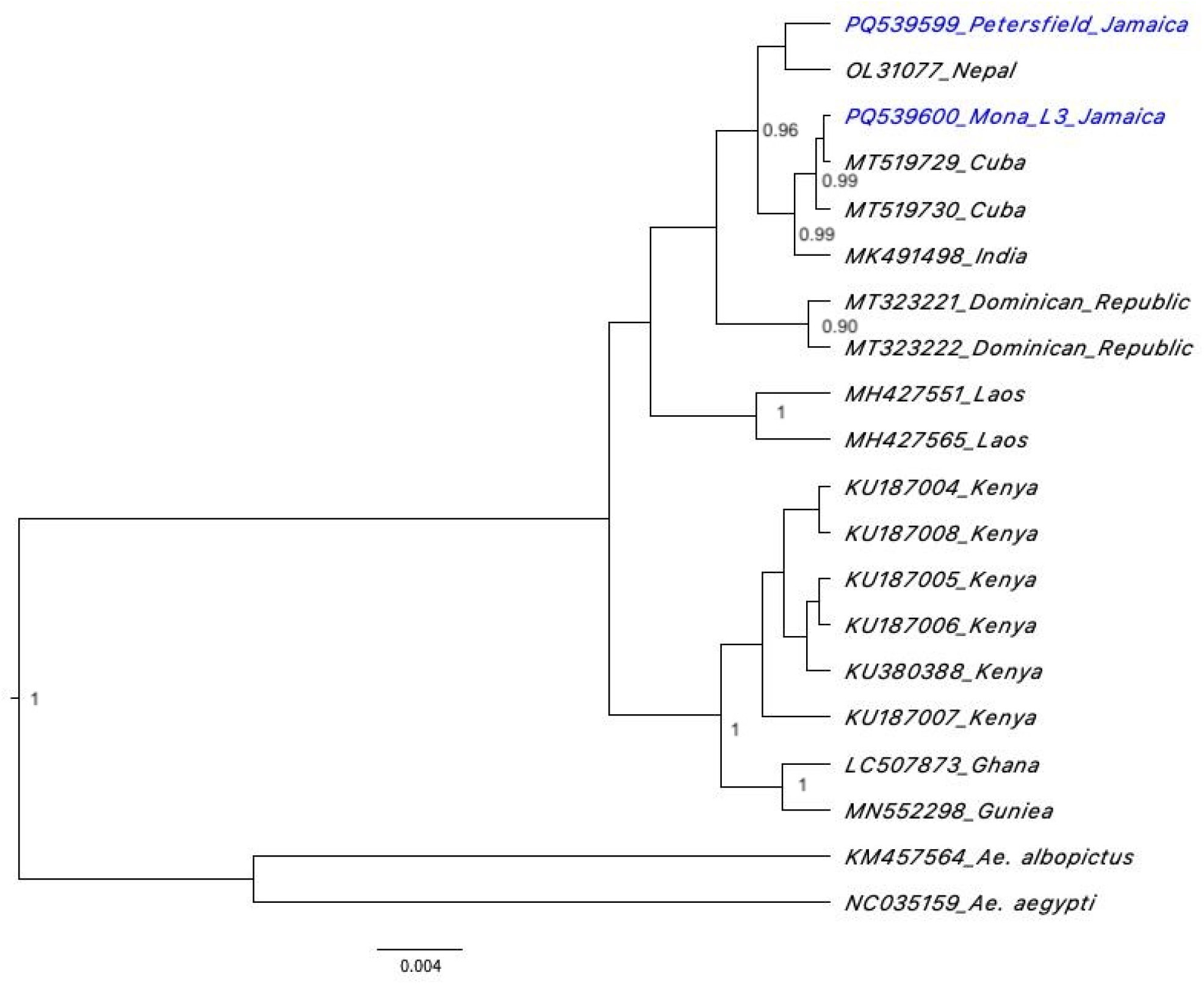
Phylogenetic tree for COI gene extracted from the mitogenomes generated in this study and available reference *Ae. vittatus* sequences. The numbers at the nodes represent posterior probabilities based on Bayesian inferences. The tree was constructed using BEAST 2 with the Tamura 3-parameter model.

## Discussion

The recent identification of invasive *Ae. vittatus* mosquito in Cuba, Dominican Republic and now in Jamaica is a significant finding with far-reaching implications for arbovirus transmission, surveillance, and vector control efforts on each island. *Aedes vittatus* was identified in four parishes representing much of the geographic expanse of Jamaica. The extensive distribution of the species across the island suggests that it may have persisted undetected for some time but appears to be present in low densities, although the collections were relatively limited.

Additionally, the discovery of larvae indicate that, *Ae. vittatus* populations are reproductively active and likely well-established. These findings highlight the potential for new dynamics in vector-borne disease transmission and the need for more extensive vector surveillance on the island. Understanding the full extent of the current distribution of this vector in Jamaica will dictate the level of control that is required.

*Aedes vittatus* an aggressive biter, is reported as a highly anthropophilic species, and was even suspected of playing a significant role in the initial transmission of YFV during the 1978–79 epidemic in The Gambia [46]. A laboratory infectivity study demonstrated *Ae. vittatus* competency in transmitting all four dengue serotypes [25]. Mavale et al. [25] indicated that although the infection rate of various dengue serotypes in *Ae. vittatus* was low, the mosquito salivary glands were infected, implying their potential to transmit or maintain the virus where dengue is endemic. A single study has also isolated dengue serotype 2 (DENV-2) from field-caught *Ae. vittatus* specimens in Senegal [47], and the vector exhibited a higher dissemination rate of DENV-2 when compared to other vectors [48]. As the timing of the introduction of *Ae vittatus* into Jamaica is currently unknown, it raises the question about the potential role of this species in past arboviral epidemics in Jamaica. In the 2018-2019 dengue epidemic in Jamaica for instance, both dengue serotypes 2 and 3 were recorded among infected patients [49].

Moreover, it is worth noting that one of the locations where the *Ae. vittatus* was collected is a residential area of Mona. Although active breeding sites were not identified at this location, this indicates the species’ ability to adapt and coexist in peri-urban/urban environments. At Walkerswood there were signs of human activity, which is significant as human encroachment into forested habitats increases potential exposure to *Ae. vittatus* and the pathogens they may transmit. *Aedes vittatus* was also collected from Myersville, St Elizabeth, a rural farming community in Jamaica. The presence of discarded refrigerators which served as human-made breeding sites for *Ae. vittatus* in this forested area underscores the impact of humans in facilitating the introduction and establishment of new species. Although anthropophilic, studies have highlighted the potential of this vector to feed on both human and animals [50, 51] which can increase the possibility of acting as a bridge vector for zoonotic pathogens. Therefore, the identification of *Ae. vittatus* in rural and peri-urban communities throughout Jamaica needs to be further evaluated. To compound these concerns even further, is the scarcity of information on arboviral transmission regarding distribution of vectors, transmission competency and host use in Jamaica and the wider Caribbean. Hence, studies which seek to highlight and report the geographical and ecological expansion and adaptation of vectors are of paramount importance.

In this study both morphological identification and molecular verification were utilized to confirm the presence of *Ae. vittatus* in Jamaica. Furthermore, we have reported the first complete mitogenomes of *Ae. vittatus* using a genome skimming approach. Sequencing of the entire mitogenome of *Ae. vittatus* provides more extensive gene coverage which can assist with more accurate molecular identification of this species and differentiation of other *Aedes* taxa. This is crucial for *Ae. vittatus* given the current misidentification of specimens as *Ae. cogilli* in GenBank which was highlighted in both the Cuban and Dominican Republic reports [13, 14]. Phylogenetic analysis conducted using a 539 bp segment of the mitochondrial COI gene may suggest multiple introductions of *Ae. vittatus* into Jamaica. Interestingly the specimen from Mona, location 3 was identical to the Cuban sample. This is suggestive of the potential role which increased trade partnerships can play in the introduction of invasive vectors. Further work will need to be undertaken to fully understand the origin(s) of the introduction of this mosquito species and dispersal throughout the country.

### Implication: Public health significance

*Aedes vittatus* poses a growing public health threat due to its extensive geographic range and proven capacity to transmit various medically important arboviruses, such as yellow fever, dengue, chikungunya, and Zika viruses [12]. Furthermore, this species exhibits a propensity to reproduce in the vicinity of human habitats and has a strong attraction to seek out human hosts [11]. The detection of this vector in the Caribbean [13-15] raises concerns regarding its possible involvement in pathogen transmission, especially in areas where established vectors such as *Ae. aegypti* and *Ae. albopictus* are controlled. The species’ adaptation to diverse natural and human-made breeding sites increases its ability to survive and proliferate in new regions. Additionally, this species has demonstrated transovarial transmission of dengue virus, a powerful potential mechanism for sustaining viral circulation between outbreaks [52]. Due to its invasive capability, enhanced by global trade and transportation, *Ae. vittatus* poses an increasing epidemiological threat in the Caribbean and Americas. Regional surveillance and vector control strategies are essential for alleviating its effects and averting future outbreaks of arbovirus-associated illnesses. Therefore, an increased and robust surveillance approach, grounded in regional communication and partnership can be used to promote awareness and the implementation of proactive control measures to prevent the potential introduction or spread of this invasive vector.

## Data Availability

The dataset generated in this study is available on the NCBI database BioProject accession PRJNA1172362. The assembled mitochondrial sequences are openly available in the NCBI GenBank database under the accession numbers PQ539599 – PQ539600.

## Acknowledgments

Simmoy A. A. Noble is a Global Infectious Diseases Scholars and received mentored research training in the development of this manuscript.

## Supporting information

**S1 Fig.**
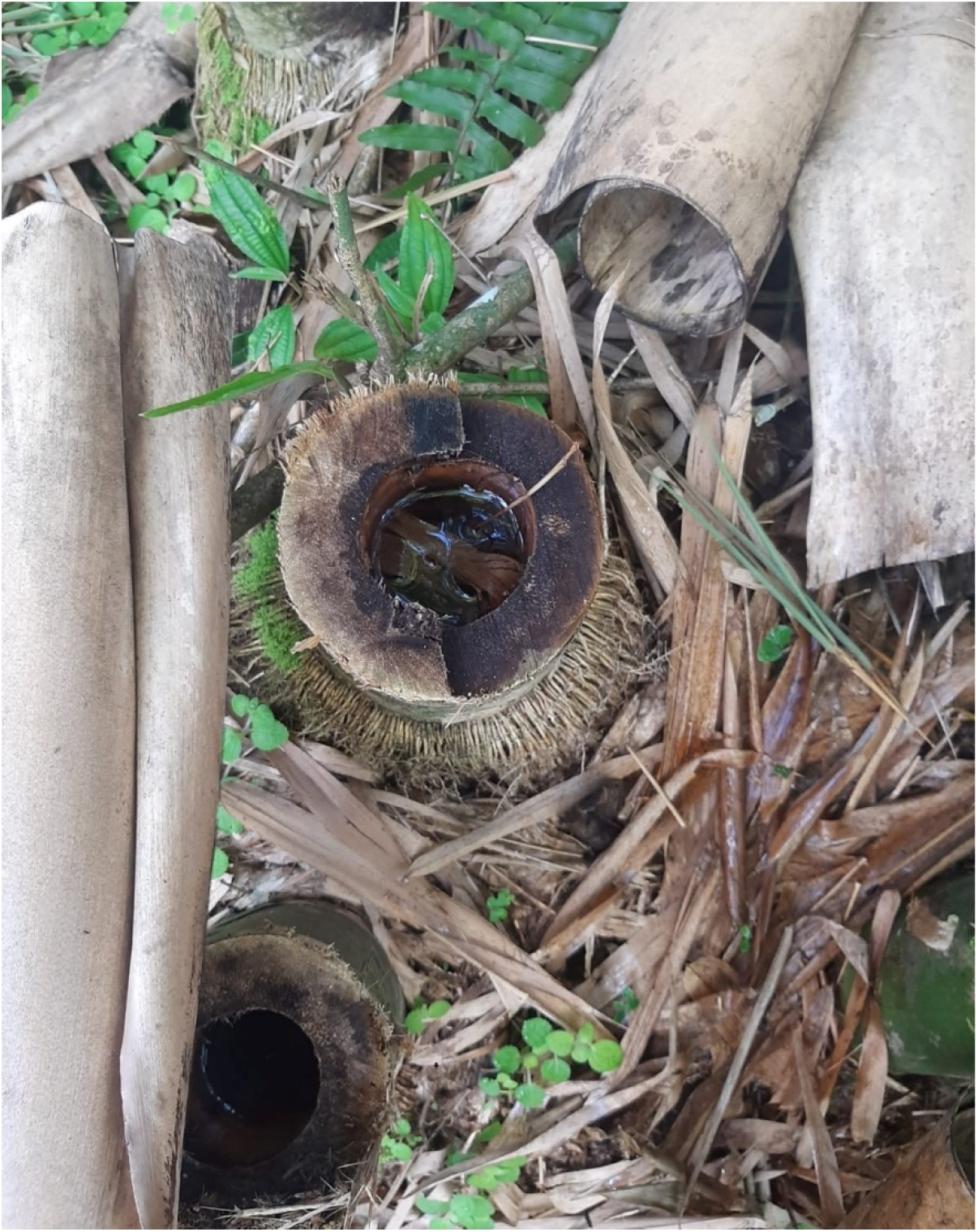
Cut bamboo indicating human activities which served as mosquito breeding site.

## References

1. Manguin S, Boëte C. Global impact of mosquito biodiversity, human vector-borne diseases and environmental change. The importance of biological interactions in the study of biodiversity. 2011:27–50. doi: 10.5772/22970.

2. Pyšek P, Hulme PE, Simberloff D, Bacher S, Blackburn TM, Carlton JT, et al. Scientists’ warning on invasive alien species. Biological Reviews. 2020;95(6):1511–34. doi: 10.1111/brv.12627.

3. Morens DM, Folkers GK, Fauci AS. Emerging infections: a perpetual challenge. The Lancet infectious diseases. 2008;8(11):710–9. doi: 10.1016/S1473-3099(08)70256-1.

4. O’meara G, Gettman A, Evans Jr L, Curtis G. The spread of Aedes albopictus in Florida. American Entomologist. 1993;39(3):163–73. doi: 10.1093/ae/39.3.163.

5. Research & Market Intelligence Unit – Jamaica Tourist Board. Overview 2021.[internet]. 2021 [2025 March 13]. Report No.

6. Research & Market Intelligence Unit – Jamaica Tourist Board. Overview 2023.[internet]. 2024 [2025 March 13]. Report No.

7. The Observatory of Economic Complexity. Jamaica/Cuba[internet]. OEC; 2023 [2025 March 13]. Available from: https://oec.world/en/profile/bilateral-country/jam/partner/cub.

8. The Observatory of Economic Complexity. `Dominica Republic/Jamaica[internet]. OEC; 2023 [2025 March 13]. Available from: https://oec.world/en/profile/bilateral-country/dom/partner/jam.

9. Benedict MQ, Levine RS, Hawley WA, Lounibos LP. Spread of the tiger: global risk of invasion by the mosquito Aedes albopictus. Vector-borne and zoonotic Diseases. 2007;7(1):76–85. doi: 10.1089/vbz.2006.0562.

10. Ali I, Mundle M, Anzinger JJ, Sandiford SL. Tiger in the sun: A report of Aedes albopictus in Jamaica. Acta tropica. 2019;199:105112. doi: 10.1016/j.actatropica.2019.105112.

11. Díez-Fernández A, Martínez-de la Puente J, Ruiz S, Gutiérrez-López R, Soriguer R, Figuerola J. Aedes vittatus in Spain: current distribution, barcoding characterization and potential role as a vector of human diseases. Parasites & vectors. 2018;11:1–6. doi: 10.1186/s13071-018-2879-4.

12. Sudeep A, Shil P. Aedes vittatus (Bigot) mosquito: An emerging threat to public health. Journal of Vector Borne Diseases. 2017;54(4):295–300. doi: 10.4103/0972-9062.225833.

13. Alarcón-Elbal PM, Rodríguez-Sosa MA, Newman B, Sutton W. The first record of Aedes vittatus (Diptera: Culicidae) in the Dominican Republic: Public health implications of a potential invasive mosquito species in the Americas. Journal of Medical Entomology. 2020;57(6):2016–21. doi: 10.1093/jme/tjaa128.

14. Pagac BB, Spring AR, Stawicki JR, Dinh TL, Lura T, Kavanaugh MD, et al. Incursion and establishment of the Old World arbovirus vector Aedes (Fredwardsius) vittatus (Bigot, 1861) in the Americas. Acta Tropica. 2021;213:105739. doi: 10.1016/j.actatropica.2020.105739.

15. Díaz-Martínez I, Diéguez-Fernández L, Santana-Águila B, de la Paz EMA, Ruiz-Domínguez D, Alarcón-Elbal PM. Nueva introducción de Aedes vittatus (diptera: Culicidae) en la región centro-oriental de cuba: Caracterización ecológica y relevancia médica. InterAmerican Journal of Medicine and Health. 2021;4. doi: doi.org/10.31005/iajmh.v4i.175.

16. Chen DH, He SL, Fu WB, Yan ZT, Hu YJ, Yuan H, et al. Mitogenome-based phylogeny of mosquitoes (Diptera: Culicidae). Insect Science. 2024;31(2):599–612. doi: 10.1111/1744-7917.13251.

17. Zé-Zé L, Borges V, Osório HC, Machado J, Gomes JP, Alves MJ. Mitogenome diversity of Aedes (Stegomyia) albopictus: Detection of multiple introduction events in Portugal. PLOS Neglected Tropical Diseases. 2020;14(9):e0008657. doi: 10.1371/journal.pntd.0008657.

18. Battaglia V, Agostini V, Moroni E, Colombo G, Lombardo G, Rambaldi Migliore N, et al. The worldwide spread of Aedes albopictus: New insights from mitogenomes. Frontiers in genetics. 2022;13:931163. doi: 10.3389/fgene.2022.931163.

19. BALATSOS G, Karathanasi V, Evangelou V, Kapantaidaki D, Tegos N, Panagopoulou A, et al. Mitogenome Diversity and Phylogenetic Insights of Aedes albopictus in Greece. bioRxiv. 2024:2024.12. 20.629447. doi: 10.1101/2024.12.20.629447.

20. Ibáñez-Justicia A, Van De Vossenberg B, Warbroek T, Teekema S, Jacobs F, Zhao T, et al. Tracking Asian tiger mosquito introductions in the Netherlands using Nextstrain. Journal of the European Mosquito Control Association. 2022;40(1):11–21. doi: 10.52004/JEMCA2021.0006.

21. Ma X-x, Wang F-f, Wu T-t, Li Y, Sun X-j, Wang C-r, et al. First description of the mitogenome and phylogeny: Aedes vexans and Ochlerotatus caspius of the Tribe Aedini (Diptera: Culicidae). Infection, Genetics and Evolution. 2022;102:105311. doi: 10.1016/j.meegid.2022.105311.

22. Seok S, Jacobsen CM, Romero-Weaver AL, Wang X, Nguyen VT, Collier TC, et al. Complete mitogenome sequence of Aedes (Hulecoeteomyia) japonicus japonicus from Hawai’i Island. Mitochondrial DNA Part B. 2023;8(1):64–8. doi: 10.1080/23802359.2022.2161328

23. Zadra N, Tatti A, Silverj A, Piccinno R, Devilliers J, Lewis C, et al. Shallow Whole-Genome Sequencing of Aedes japonicus and Aedes koreicus from Italy and an Updated Picture of Their Evolution Based on Mitogenomics and Barcoding. Insects. 2023;14(12):904. doi: 10.3390/insects14120904.

24. e Silva LHdS, da Silva FS, de Almeida Medeiros DB, Cruz ACR, da Silva SP, de Oliveira Aragão A, et al. Description of the mitogenome and phylogeny of Aedes spp.(Diptera: Culicidae) from the Amazon region. Acta Tropica. 2022;232:106500. doi: 10.1016/j.actatropica.2022.106500.

25. Mavale M, Ilkal M, Dhanda V. Experimental studies on the susceptibility of Aedes vittatus to dengue viruses. Acta virologica. 1992;36(4):412-6. PubMed Central PMCID: PMC 1362325.

26. Jupp P, McIntosh B. Aedes furcifer and other mosquitoes as vectors of chikungunya virus at Mica, northeastern Transvaal, South Africa. J Am Mosq Control Assoc. 1990;6(3):415-20. PubMed Central PMCID: PMC1977875.

27. Mourya D, Banerjee K. Experimental transmission of Chikungunya virus by Aedes vittatus mosquitoes. 1987. PubMed Central PMCID: PMC3428961.

28. Diallo D, Sall AA, Diagne CT, Faye O, Faye O, Ba Y, et al. Zika virus emergence in mosquitoes in southeastern Senegal, 2011. PloS one. 2014;9(10):e109442. doi: 10.1371/journal.pone.0109442.

29. Sudeep A, Mohandas S, Bhanarkar S, Ghodke Y, Sonawane P. Vector competence of Aedes vittatus (Bigot) mosquitoes from India for Japanese encephalitis, West Nile, Chandipura and Chittoor viruses. Journal of Vector Borne Diseases. 2020;57(3):234–9. doi: 10.4103/0972-9062.311776.

30. Irving-Bell R, Inyang E, Tamu G. Survival of Aedes vittatus (Diptera: Culicidae) eggs in hot, dry rockpools. Tropical medicine and parasitology: official organ of Deutsche Tropenmedizinische Gesellschaft and of Deutsche Gesellschaft fur Technische Zusammenarbeit (GTZ). 1991;42(1):63-6. doi: 2052860.

31. Service M. Studies on the biology and taxonomy of Aedes (Stegomyia) vittatus (Bigot)(Diptera: Culicidae) in northern Nigeria. 1970. doi: 10.1111/j.1365-2311.1970.tb00529.x.

32. Tewari S, Thenmozhi V, Katholi C, Manavalan R, Munirathinam A, Gajanana A. Dengue vector prevalence and virus infection in a rural area in south India. Tropical Medicine & International Health. 2004;9(4):499–507. doi: 10.1111/j.1365-3156.2004.01103.x.

33. Diallo D, Diagne CT, Hanley KA, Sall AA, Buenemann M, Ba Y, et al. Larval ecology of mosquitoes in sylvatic arbovirus foci in southeastern Senegal. Parasites & vectors. 2012;5:1–17. doi: 10.1186/1756-3305-5-286.

34. Mondal R, Devi NP, Bhattacharya S. Seasonal prevalence and host preference of some medically important Aedes species of Doon Valley, India. 2021. doi: 10.24321/0019.5138.202144.

35. Kumari R, Kumar K, Chauhan LS. First dengue virus detection in Aedes albopictus from Delhi, India: Its breeding ecology and role in dengue transmission. Tropical Medicine & International Health. 2011;16(8):949–54. doi: 10.1111/j.1365-3156.2011.02789.x.

36. Belkin JN, Heinemann SJ, Page WA. The culicidae of Jamaica 1970. 1-458]. Available from: https://mosquito-taxonomic-inventory.myspecies.info/sites/mosquito-taxonomic-inventory.info/files/Belkin%20et%20al%201970.pdf.

37. Walter Reed Biosystematics Unit. Aedes vittatus species page [internet]. Walter Reed Biosystematics Unit Website WRBU; 2021 [cited 2025]. Available from: http://wrbu.si.edu/vectorspecies/mosquitoes/ae_vittatus.

38. Chen T-Y, Vorsino AE, Kosinski KJ, Romero-Weaver AL, Buckner EA, Chiu JC, et al. A magnetic-bead-based mosquito DNA extraction protocol for next-generation sequencing. Journal of Visualized Experiments (JoVE). 2021;(170):e62354. doi: 10.3791/62354.

39. Sandiford SL, Noble SA, Pierre SA, Norris DE, Ali R. The first mitochondrial genome of Haemagogus equinus from Jamaica. F1000Research. 2024;13:1504. doi: 10.12688/f1000research.159115.1.

40. Dierckxsens N, Mardulyn P, Smits G. NOVOPlasty: de novo assembly of organelle genomes from whole genome data. Nucleic Acids Research. 2016;45(4):e18–e. doi: 10.1093/nar/gkw955.

41. Bernt M, Donath A, Jühling F, Externbrink F, Florentz C, Fritzsch G, et al. MITOS: improved de novo metazoan mitochondrial genome annotation. Molecular phylogenetics and evolution. 2013;69(2):313–9. doi: 10.1016/j.ympev.2012.08.023.

42. Price G NA, Grüning BA, Schatz MC. The Galaxy platform for accessible, reproducible, and collaborative data analyses: 2024 update. Nucleic acids research. 2024;52(W1):W83–W94. doi: 10.1093/nar/gkae410.

43. Posada D. jModelTest: phylogenetic model averaging. Molecular biology and evolution. 2008;25(7):1253–6. doi: 10.1093/molbev/msn083.

44. Bouckaert R, Heled J, Kühnert D, Vaughan T, Wu C-H, Xie D, et al. BEAST 2: a software platform for Bayesian evolutionary analysis. PLoS computational biology. 2014;10(4):e1003537. doi: 10.1371/journal.pcbi.1003537.

45. Hartke J, Reuss F, Kramer IM, Magdeburg A, Deblauwe I, Tuladhar R, et al. A barcoding pipeline for mosquito surveillance in Nepal, a biodiverse dengue-endemic country. Parasites & vectors. 2022;15(1):145.

46. Germain M, Monath P, Bryan J, Salaun J, Renaudet J. Aspects and epidemiological correlations Am J Trop Med Hyg. 1980;29(5):929–40.

47. Diallo M, Ba Y, Sall AA, Diop OM, Ndione JA, Mondo M, et al. Amplification of the sylvatic cycle of dengue virus type 2, Senegal, 1999–2000: entomologic findings and epidemiologic considerations. Emerging infectious diseases. 2003;9(3):362. doi: 10.3201/eid0903.020219.

48. Diallo M, Sall AA, Moncayo AC, Ba Y, Fernandez Z, Ortiz D, et al. Potential role of sylvatic and domestic African mosquito species in dengue emergence. The American journal of tropical medicine and hygiene. 2005;73(2):445-9. PubMed Central PMCID: PMC16103619.

49. Lue AM, Richards-Dawson M-AEH, Gordon-Strachan GM, Kodilinye SM, Dunkley-Thompson JAT, James-Powell TD, et al. Severity and outcomes of Dengue in hospitalized Jamaican children in 2018–2019 during an epidemic surge in the Americas. Frontiers in medicine. 2022;9:889998. doi: 10.3389/fmed.2022.889998.

50. Service M. The Identification of Blood-Meals from Culicine Mosquitos from Northern Nigeria. 1965. doi: doi:10.1017/S0007485300049749.

51. Wilson JJ, Sevarkodiyone S. Host preference of blood feeding mosquitoes in rural areas of southern Tamil Nadu, India. Acad J Entomol. 2015;8(2):80–3. doi: 10.5829/idosi.aje.2015.8.2.94106.

52. Angel B, Sharma K, Joshi V. Association of ovarian proteins with transovarial transmission of dengue viruses by Aedes mosquitoes in Rajasthan, India. Indian Journal of Medical Research. 2008;128(3):320-3. PubMed Central PMCID: PMC19052346.

